# Optimal positioning and size of high-density electrocorticography grids for speech brain-computer interfaces

**DOI:** 10.1101/2025.03.13.643127

**Authors:** Elena C Offenberg, Julia Berezutskaya, Lennart Müller, Zachary V Freudenburg, Nick F Ramsey, Mariska J Vansteensel

## Abstract

Speech-based brain-computer interfaces (BCIs) can offer an intuitive means of communication for those who have lost the ability to speak due to paralysis. Significant progress has been made in classifying individual words from high numbers of electrocorticographic (ECoG) electrodes on the sensorimotor cortex (SMC). As implantations of larger grids with more ECoG electrodes are associated with higher surgical risk, we here examined whether confined electrode configurations can match the classification accuracy of larger grids. To this end, we analyzed data from eight able-bodied participants with high-density ECoG grids (64 to 128 electrodes) who performed a task that involved speaking 12 Dutch words. Word pronunciation was associated with changes in high frequency band activity in two SMC foci, one in the ventral SMC and another in the dorsal SMC. Using a combinatorics approach, we found that a smaller, rectangular, configuration with a surface area of 325 mm^2^ to 561 mm^2^ (32 electrodes) could achieve a word classification accuracy similar to that of the larger grids: 76±16% versus 75±17% across participants, respectively (practical chance level 16.7%). The best configurations were oriented vertically and centered on the central sulcus. These findings indicate that a 32-electrode ECoG grid placed optimally can be sufficient for achieving high word classification accuracy on a closed set of words. We conclude that targeted placement of small ECoG grids can reduce surgical demands on end users and justify energy- and complexity-efficient designs of fully implantable BCI devices for individuals with severe paralysis.

## 1. Introduction

Neurodegenerative diseases such as amyotrophic lateral sclerosis (ALS) and conditions such as brainstem stroke can cause profound motor impairments. In the most severe cases, individuals may experience complete paralysis and loss of speech (anarthria) while remaining cognitively intact—a condition known as locked-in syndrome (American Congress Of Rehabilitation Medicine, 1995, Laureys et al., 2005). The accompanying inability to verbally communicate compromises the quality of life of the affected individuals (Felgoise et al., 2016). Research demonstrates that employing augmentative and assistive communication (AAC) devices can improve the quality of life of people with severe motor impairments (Körner et al., 2013, Rousseau et al., 2015). Conventional AAC devices, however, rely on residual movement, which not all affected individuals may be able to reliably produce, resulting in limited utility (Spataro et al., 2014, Kageyama et al., 2020).

Brain-Computer Interfaces (BCIs) that record signals directly from the brain can offer an alternative communication solution for people with motor impairments. Particularly promising are implantable BCIs using subdural electrocorticography (ECoG) electrodes, which offer high spatial and temporal resolution and high signal-to-noise ratio (Ball et al., 2009). Furthermore, implanted ECoG systems have shown signal stability and safety over several years after implantation, with minimal need for recalibration (Larzabal et al., 2021, Nair et al., 2020, Wyse-Sookoo et al., 2024, Vansteensel et al., 2024).

One intuitive way of controlling a communication-BCI is by attempting to speak, which is among the strategies preferred by individuals with locked-in syndrome (Branco et al., 2021). Recent years have seen large strides in speech decoding using ECoG signals from SMC. Research in able-bodied participants has demonstrated reliable classification of individual words from grids with 10 mm electrode spacing (Martin et al., 2016), 3-4 mm spacing (Berezutskaya et al., 2023) and 1 mm spacing (Kellis et al., 2010), referred to as clinical, High-Density (HD) and micro ECoG respectively. These results, particularly using HD-ECoG grids, have already been successfully applied in people with severe paralysis in research settings. Moses and colleagues used HD-ECoG to classify 50 distinct words in a person unable to speak due to a brainstem stroke, with a median word error rate of 25.6% (Moses et al., 2021). In another individual with a brainstem stroke, accurate classification of a much larger vocabulary was made possible with the aid of language models, resulting in a median word error rate of 25% (Metzger et al., 2023). Luo and colleagues reached a median accuracy of 90.59% on six intuitive speech commands over a period of several months in a person with severe dysarthria due to ALS (Luo et al., 2023).

Next to demonstrating remarkable results of classifying words using ECoG BCIs in people with severe motor impairment, recent studies appear to be setting a trend for using an increasing number of ECoG electrodes. Earlier BCI studies restoring communication in people with paralysis used 8 to 16 contacts (Vansteensel et al., 2016, Oxley et al., 2021), whereas later studies report on grids with 128 to 253 electrodes (Moses et al., 2021, Metzger et al., 2023, Angrick et al., 2024). While this trend is partly driven by an increase in electrode density, the size of implanted grids has also expanded. However, larger grids have two disadvantages. First, they lead to increased surgical risks and burden for the user by necessitating a large craniotomy, compared to smaller grids, which could be inserted via burr holes. Second, grids with higher electrode counts require increasingly complex electronics for fully implantable devices, posing significant technical challenges regarding size, power efficiency and temperature control.

In practice, only a subset of electrodes in large coverages contributes the vast majority of information for word classification (Moses et al., 2021, Metzger et al., 2023, Angrick et al., 2024). The perceived need for extensive electrode coverage may result partly from the lack of knowledge about which areas of the SMC are most informative for speech decoding. While large grids may be appropriate for research and for the proof of concept phase, the translation towards viable clinical applications necessitates, among other things, minimizing the risk for end users to the largest extent possible. Answering open questions such as whether larger and denser grids are necessary, how many ECoG electrodes are needed for different levels of speech-BCI functionalities, and where electrodes should be positioned to be most informative will help the development of safer BCI devices.

In this study, we investigated whether ECoG grids of a limited size could yield high classification performance and assessed where such grids would best be placed. We focused on classifying twelve individual words in eight able-bodied human participants with temporarily implanted 64-to 128-electrode HD-ECoG grids. We found two SMC foci that were the most active during word pronunciation, one in the dorsal and one in the ventral SMC. We then trained word classifiers to identify the optimal number and location of electrodes needed for best classification accuracy. We found that across participants, a 32-electrode configuration covering a cortical surface area of 325 mm^2^ to 561 mm^2^, yielded a classification accuracy comparable to that of the full grid with areas of 1,225 mm^2^ to 2,806 mm^2^. The optimally positioned 32-electrode configurations were oriented vertically and covered the central sulcus.

## 2. Methods

### 2.1 Data collection

Eight able-bodied participants between the ages of 19 and 51, 4 female, were temporarily implanted with HD-ECoG grids over their SMC. All grids were placed on the left hemisphere with the exception of P2 (Figure 1). In all participants, the HD grids were placed on healthy brain tissue for the purpose of research. Surgery was dictated solely by clinical considerations, and HD grids were only placed if the SMC could be reached safely and the participant had consented. Seven participants (P1-P7) were epilepsy patients, and ECoG data was recorded while they were being monitored in preparation for epilepsy resection surgery. For the final participant P8, the HD-ECoG grid was placed temporarily on the SMC during awake tumor resection surgery, and ECoG data was recorded intraoperatively. The research was approved by the Medical Ethical Committee of the University Medical Center Utrecht in accordance with the Declaration of Helsinki (2024). All participants gave written informed consent to participate in this research.

**Figure 1.**
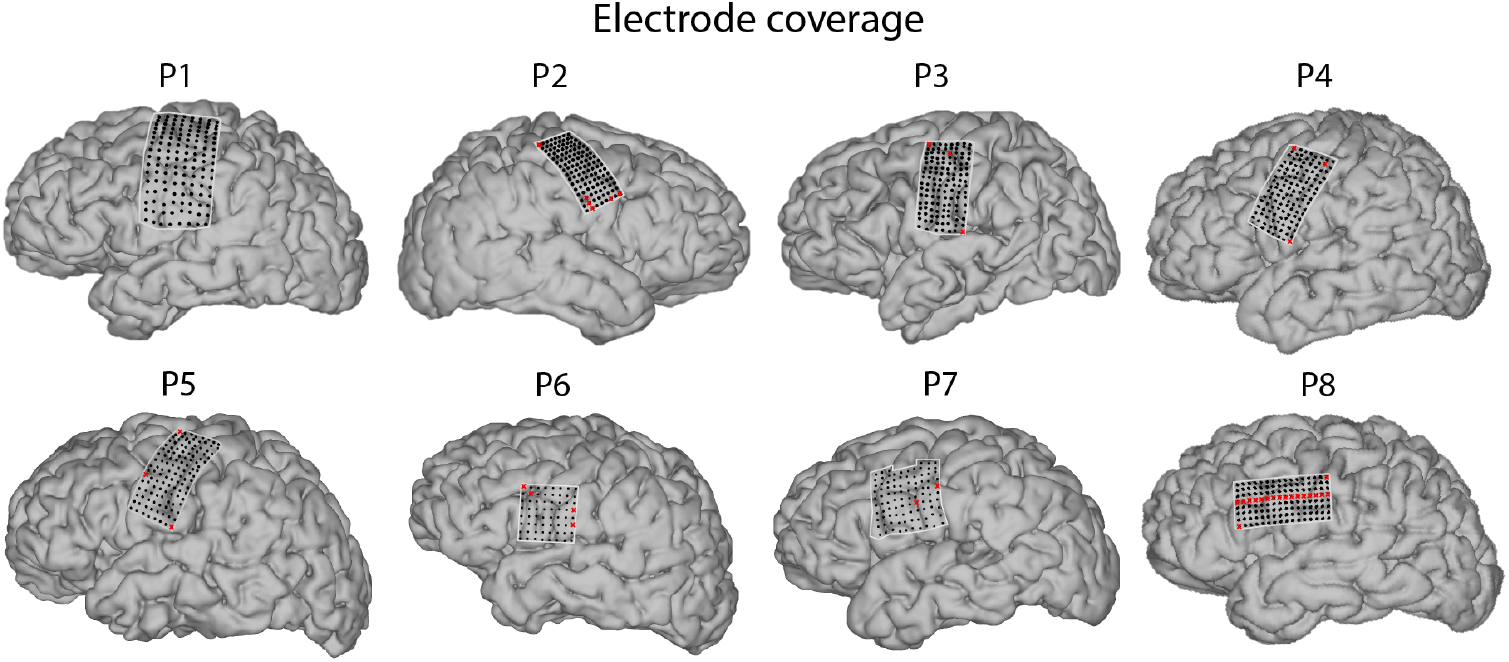
Placement of HD-ECoG grids on the sensorimotor cortex for all eight participants. Electrodes removed from the analyses for poor signal quality are marked in red.

Six of the eight participants (P1-P5, P8) were implanted with 128-electrode HD-ECoG grids, one (P6) with a 64-electrode HD-ECoG grid and one (P7) with a 96-electrode HD-ECoG grid. The data from participants P1, P2, P3, P5, and P8 was previously reported on as part of an earlier study with a different objective (Berezutskaya et al., 2023). The technical details for the grids are listed in Table 1. In all cases, neural data was recorded using a Blackrock neural signal processor system at 2 kHz (Blackrock Microsystems). Microphone recordings of the participants’ voice were acquired simultaneously at 30 kHz and synchronized with the brain signals.

**Table 1.**
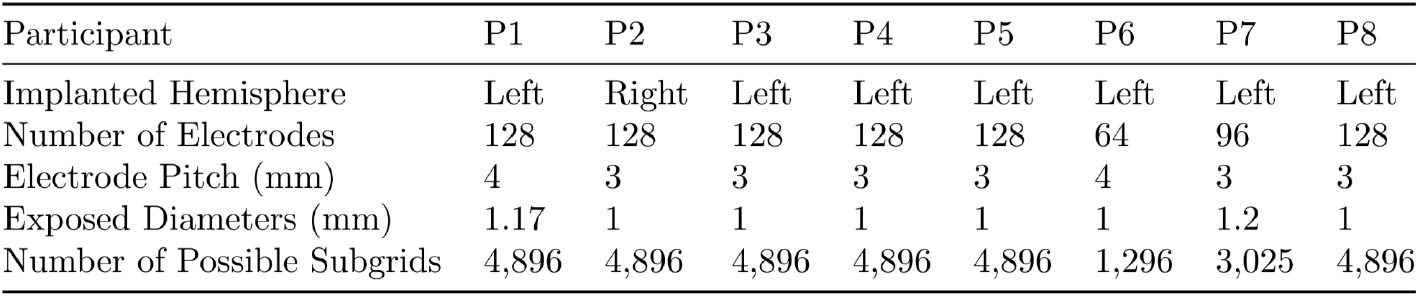
Information about HD-ECoG grids for each participant, including technical details as well as the number of possible subgrids in the exhaustive grid search as described in Section 2.6.

| Participant | P1 | P2 | P3 | P4 | P5 | P6 | P7 | P8 |
| --- | --- | --- | --- | --- | --- | --- | --- | --- |
| Implanted Hemisphere | Left | Right | Left | Left | Left | Left | Left | Left |
| Number of Electrodes | 128 | 128 | 128 | 128 | 128 | 64 | 96 | 128 |
| Electrode Pitch (mm) | 4 | 3 | 3 | 3 | 3 | 4 | 3 | 3 |
| Exposed Diameters (mm) | 1.17 | 1 | 1 | 1 | 1 | 1 | 1.2 | 1 |
| Number of Possible Subgrids | 4,896 | 4,896 | 4,896 | 4,896 | 4,896 | 1,296 | 3,025 | 4,896 |

### 2.2 Task and data preprocessing

Participants took part in a word repetition task during which they were shown isolated words on a screen one at a time and were instructed to pronounce each word aloud (Figure 2). There were twelve unique Dutch words in the task: “allemaal”, “boe”, “ga”, “grootmoeder”, “hoed”, “ik”, “janneke”, “jip”, “kom”, “plukken”, “spelen”, and “waar”. These words, as well as “-”, which represented rest trials, were presented on screen ten times each in random order. For all participants except P8, each word cue was shown for 1.5 seconds, followed by an inter-trial “+”-cue shown for 2.5 seconds until the next word was presented. The participants were instructed to pronounce the words upon seeing the word cue. Due to time constraints in the intraoperative setting, P8 was only shown each word for 1 second and the inter-trial screen for 2 seconds, and was not presented with any rest trials. Since the task performed by P1-P7 only included 10 rest trials as opposed to 120 word trials (10 repetitions of 12 unique words) and because there were no rest trials in the task of P8, rest trials were not used for further analyses. Instead, silence fragments from between word pronunciations were identified and used for all participants to compare with active (speech pronunciation) trials in order to test for task-specific activation. In addition, some words were mispronounced or repeated during a single trial, causing the number of repetitions per word to vary between 7 and 12 for P1, P3, P7 and P8, with P2, P4, P5 and P6 providing exactly 10 repetitions for each word.

**Figure 2.**
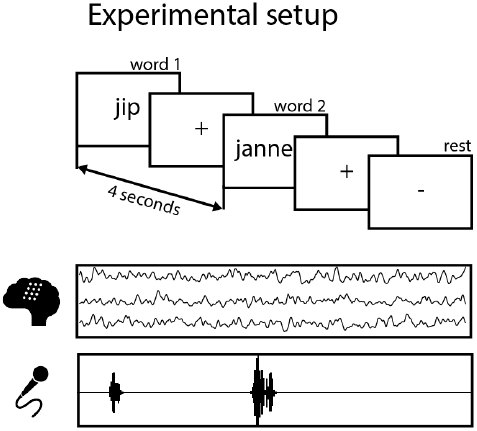
The experimental setup of the word repetition task, adapted from (Berezutskaya et al., 2023).

The recorded HD-ECoG data was preprocessed following a standard procedure. First, electrodes displaying noisy or flat signals, as determined through visual inspection, were excluded from further analysis. Additionally, in grids with upwards-facing reference electrodes (typically, electrodes #1 and #128), we excluded such reference electrodes, leading to the following electrode counts used in all analyses: P1 – 128, P2 – 122, P3 – 125, P4 – 125, P5 – 125, P6 – 60, P7 – 94 and P8 – 110 electrodes. A notch filter for line noise was applied to the brain signals at 50 Hz and harmonics. After common average referencing the signal of included electrodes, Gabor wavelet decomposition was used to extract band-pass power signals of different frequency bands in 1 Hz bins. Per frequency band, these power signals were then averaged across the 1 Hz bins (delta [0,5 – 3 Hz], theta [4 – 7 Hz], alpha [8 – 12 Hz], beta [13 – 30 Hz], and the high frequency band (HFB) [70 – 170 Hz]), log-transformed and downsampled to 50 Hz. Per participant, the mean power in each frequency band of each electrode *s* was normalized using the log-transform formula (Mercier et al., 2022):

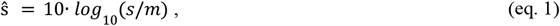

where *m* is the mean of a 5-second baseline period. The baseline period was identified per participant as a period at the start of the task where the participant was neither speaking nor being spoken to, as determined based on the synchronized microphone recordings. This log-transform normalization procedure was selected to correct for the 1/f signal fall-off, minimizing the differences in signal distributions across frequency bands.

Word on- and offsets were identified based on microphone recordings synchronized with the brain data. For participants P1-P4, P6, and P8, word on- and offsets were determined manually. To expedite the process, for P5 and P7, the Montreal Forced Aligner tool with the pretrained Dutch acoustic model was used for automatic voice onset and offset detection (McAuliffe et al., 2017), followed by a manual check.

### 2.3 Task-specific activation

To test for task-specific activation per electrode and frequency band, a two-tailed t-test comparing the means of brain activity during speech and silence fragments was performed. Speech fragments were defined as the signals from voice onset to offset. Since the longest word pronunciation across participants was 0.975 seconds, silence fragments were defined as the periods starting from 1 s after voice onset to 1s before the next voice onset. Any silence fragments that were shorter than 0.1s were ignored. This resulted in a mean length of speech fragments of 0.52±0.06s, and a mean length of silence fragments of 2.36±1.18s. The significance threshold was Bonferroni-corrected and set to 0.05/(N_e_⋅N_f_ ⋅N_s_), where N_e_, N_f_, and N_s_ corresponded to the number of electrodes, the number of frequency bands and the number of participants, respectively.

### 2.4 Projection into common space

In order to combine findings from all participants for group analysis and display, we applied two distinct projection methods, both relying on the same participant-specific electrode locations. For P1-P7, these locations were determined through post-implantation computed tomography scans that were coregistered to their presurgical anatomical MRI scans using FreeSurfer (Hermes et al., 2010, Fischl, 2012, Branco et al., 2018a). In the case of P8, the exact location of the grid electrodes was determined using the Gridloc method (Branco et al., 2018b).

For the first projection method, designed for visualizing results on the common brain surface, we projected all electrode positions to Montreal Neurological Institute (MNI) space using patient-specific affine transformation matrices obtained with SPM8 (Wellcome Centre for Human Neuroimaging, University College London). As a final step of the visualization, a 2D Gaussian kernel with a FWHM of 10 mm was applied to the brain surface coordinates at each electrode’s center, so that the obtained activity values faded outwards from the center of the electrode. To visualize the observed activity in relation to the parcellation from the Human Connectome Project, we utilized the Connectome Workbench software to map the regions of interest (55b and 6v) onto the MNI brain (Marcus et al., 2011, Glasser et al., 2016).

The second method was used for a quantitative analysis of task response localizations within the sensorimotor cortex. For this, we clustered the locations of t-test values across electrodes in a sensorimotor space common across participants. When multiple electrodes from different participants were mapped to the same coordinate in this shared space, their t-test values were averaged. To make this common space, we applied the Cgrid (Cartesian geometric representation with isometric dimension) method (Bruurmijn et al., 2020). Using Cgrid, the vertices of the precentral and postcentral gyrus of each individual FreeSurfer surface reconstruction were warped into a common Cartesian grid. The t-values were mapped to this Cgrid through linkage of the electrode coordinates to the closest vertices in the pial surface, then resampled to the 2D regular Cgrid, and smoothed with a 2D Gaussian kernel with a FWHM of 8 Cgrid-pixels (corresponding to approximately 10.4 mm). Subsequently, we performed connected component clustering by determining the number of two-dimensional four-connected neighborhood components (i.e. clusters) separately for precentral and postcentral gyri. For statistical testing, the t-values of the electrodes were randomly permuted 10,000 times across the electrode locations. The same analysis was then repeated, counting how often the same number of clusters appeared relative to the total permutations. A significance threshold of p = 0.05 was applied to determine statistical relevance.

### 2.5 Word classification

Support vector machines (SVMs) with linear kernels, as implemented in scikit-learn (Pedregosa et al., 2018), were utilized for the word classification. The input was brain data from a window starting at voice onset, with a participant-specific window length. This length was set to be the maximum pronunciation time (i.e., time between voice onset and offset) of that participant across all words and repetitions. Since SVMs do not support multi-class classification natively, a one-vs-one strategy was used for discriminating the 12 words. Each participant-specific dataset was divided into training and test data, ensuring that one repetition of each word was included in the test set, following the leave-one-group-out scheme. In a nested cross-validation procedure, another repetition of each word was used as a validation set in order to optimize the regularization parameter of the SVM. This was achieved using the automatic hyperparameter selection library Optuna (Akiba et al., 2019). From a potential parameter range of 10^−5^ to 10^6^, the algorithm employed a tree-structured Parzen estimator to identify the best regularization parameter within 30 steps, based on the information from preceding steps.

Accuracy, defined as the proportion of correctly classified test set words, was used as the evaluation metric. In all cases, median accuracy plus/minus the median absolute deviation is reported. To account for the limited dataset size, a binomial cumulative distribution was used to determine the statistical significance threshold of classification accuracy (Combrisson & Jerbi, 2015), resulting in 16.7% for *p* < 10^−3^.

Word classification was first applied to the full grids. We then performed a subgrid analysis, retraining the classifier on smaller configurations (see Section 2.6). Finally, recursive feature elimination was performed to identify the most informative electrodes for word classification and evaluate whether those responsive to speech were also essential for differentiating words (see Section 2.7). To compare the performances of these analyses, Wilcoxon-signed rank tests were used (Wilcoxon, 1945), with Bonferroni-correction for multiple comparisons where applicable.

### 2.6 Subgrid analysis

For the subgrid analysis, an exhaustive grid search was initially conducted. All possible rectangular subgrids for each electrode grid were determined, from the smallest possible subgrid of just one electrode (1×1) to the full grid (8×8, 10×10 or 8×16, depending on the size of the full grid). Due to differences in grid sizes, the number of possible subgrids varied for each participant and is listed in Table 1. SVM word classifiers were trained on all subgrids without hyperparameter optimization to increase efficiency. If an electrode excluded during preprocessing was part of a subgrid, it was also removed from input to the SVM subgrid classifier.

Based on these initial analyses, we chose the smallest subgrid size where added electrodes marginally improved accuracy for in-depth analysis. We then retrained the SVM classifiers using hyperparameter optimization to directly compare the performance of these smaller subgrids with that of full grids. Due to the cross-validation strategy used for training, each of the 10 folds could yield a different best-performing subgrid based on classification performance.

Since data preprocessing included common average referencing, we additionally verified whether the results were affected by using smaller subgrids. For all participants, we retrained the SVM classifiers with the smaller subgrids, this time recalculating the common average signal using only the electrodes within each subgrid as part of a full preprocessing rerun. We then compared classification accuracy between the original and reprocessed data using a Wilcoxon signed-rank test to determine whether rerunning preprocessing had any significant effect.

To calculate the surface area *a*_g_ of a subgrid *g* with *N*_w_ electrodes along the width and *N*_h_ along the height-dimension of the grid, as well as an interelectrode distance *d*_ie_ and an electrode radius *r*, the following formula was used:

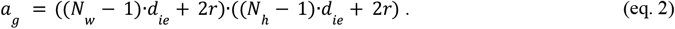

The code used for this analysis is publicly available in our BCI-sift toolbox (github.com/UMCU-RIBS/BCIsift).

### 2.7 Recursive feature elimination

For the recursive feature elimination (RFE) analysis, each individual electrode was considered to be a separate feature. RFE iteratively retrained the SVM classifier, removing the least informative feature at each step. The feature considered least informative was the electrode with the lowest average absolute weight assigned by the trained SVM. This process was repeated until only one feature remained. To be consistent with the subgrid analysis, we then selected the RFE step that used at most as many channels as the smallest subgrid size where added electrodes marginally improved accuracy in Section 2.6, and that achieved the highest accuracy on the validation set. The reported accuracy was calculated by applying the SVM trained on the selected features to the test set in a leave-one-group-out cross-validation scheme.

The code used for this analysis is publicly available in our BCI-sift toolbox (github.com/UMCU-RIBS/BCIsift).

## 3. Results

### 3.1 Task-specific activation

First, we analyzed the differences in band power between speech and silence across frequency bands and electrodes, so as to determine per frequency band which electrodes responded to speech. Across participants, HFB signals showed a significant increase in activity during speech, while the beta band typically showed a significant decrease (Figure 3). There was generally a less prominent decrease in activity in the alpha band and theta band. In the delta band, no significant differences between speech and silence were observed. Subsequent analyses excluded the less responsive frequency bands, focusing on only the HFB and beta bands. Two participants (P4 and P5) exhibited increases in the beta band during speech in several electrodes. Some participants also exhibited significant HFB decreases in a few electrodes, although most significant electrodes in this band showed an increase. To maintain consistency across all participants, all significant electrodes—whether showing increases or decreases in HFB and beta power—were included in the analysis moving forward.

**Figure 3.**
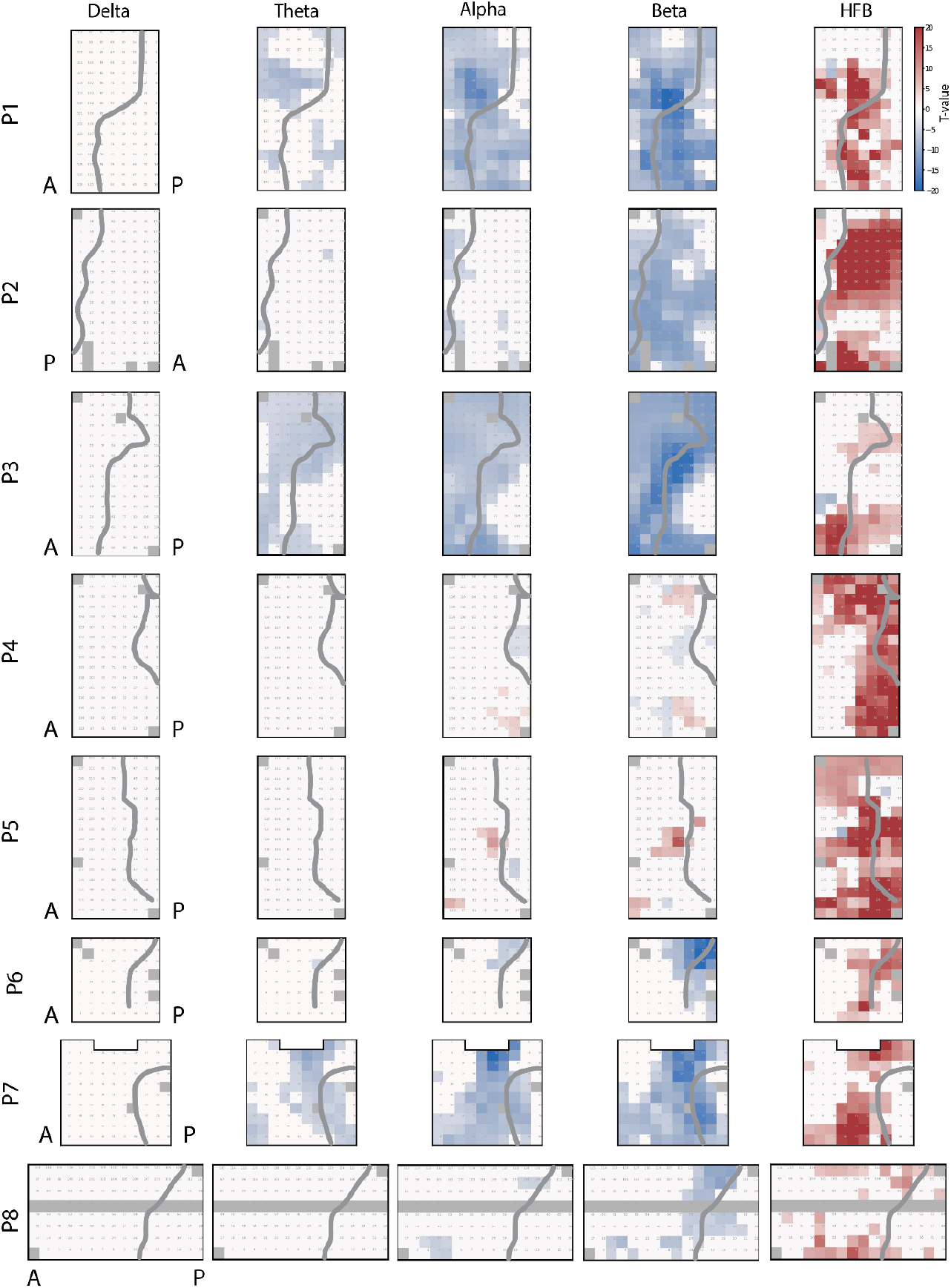
Electrodes with a significant difference in mean band power during speech and silence, as determined by t-tests. The significance threshold for the t-values varied by participant, averaging an absolute value of 4.2±0.1. Red represents the band power increasing during speech compared to rest, while blue represents the band power decreasing during speech. Electrodes in dark gray were removed from all analyses due to noisy signals or being reference electrodes. The central sulci are drawn in dark gray. All grids are located on the left hemisphere, with the exception of P2 (right hemisphere), as shown by the labels “A” for anterior and “P” for posterior direction.

The HFB activity appeared to concentrate in two distinct areas in the sensorimotor cortex, one located more dorsally and one more ventrally, while the beta decrease across participants occurred throughout all parts of SMC that were covered by electrodes (Figure 4, Figure S2). To determine whether this separation into two foci of HFB activity was significant, the connected component clustering analysis was performed separately for the precentral and postcentral gyri, as these regions are functionally and anatomically distinct. In the precentral gyrus, two distinct clusters were identified, a result that reached statistical significance through random permutation testing (p = 0.01). Significance was not reached in the postcentral gyrus.

**Figure 4.**
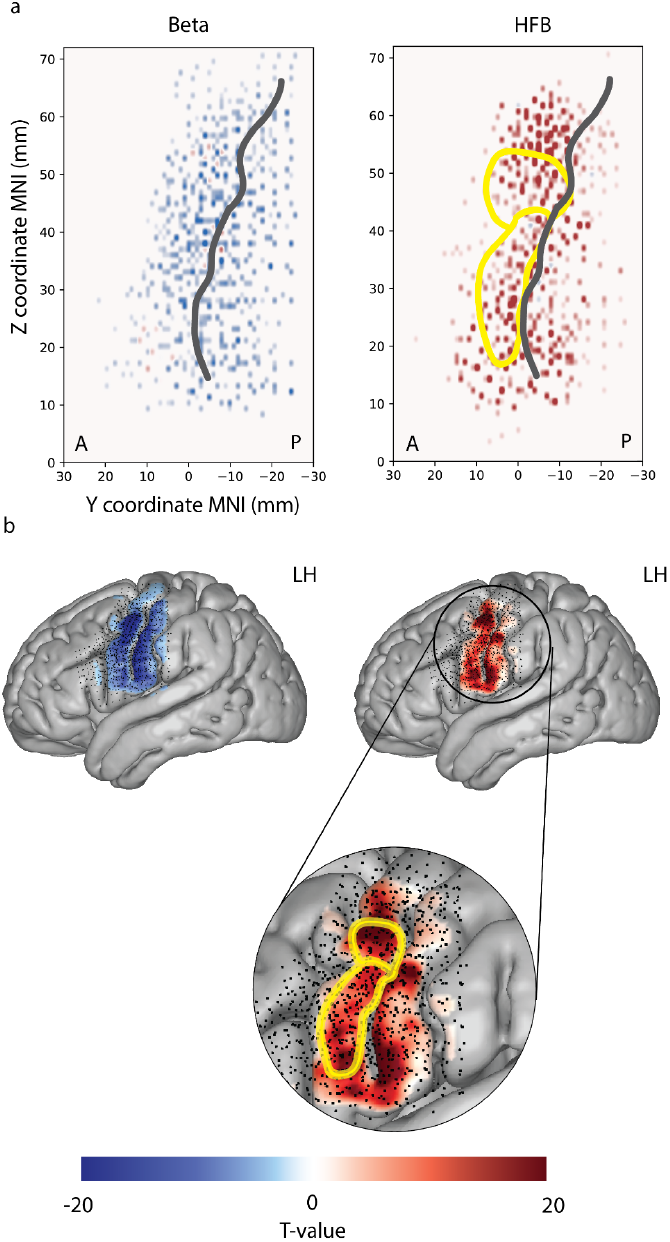
Electrodes with significant HFB and beta power responses for words vs. rest, combined for all participants in MNI space. Red indicates increased activity during speech, blue indicates decreased activity, and darker shades represent stronger t-test divergences between speech and silence. a) Significant electrodes in a 2D MNI projection (x-coordinate removed) with anterior (A) and posterior (P) directions labeled. b) Significant electrodes mapped onto the MNI brain, with black dots representing all electrodes and colors showing smoothed t-values (see Methods). For visualization purposes, data from P2 is mapped onto the left hemisphere (LH). The areas 55b and 6v, which are known to be integral to speech processing and production, are highlighted in yellow on the right parts of both subplots, with a zoom-in on the relevant area in b).

### 3.2 Full grid analysis

Median word classification accuracy achieved with full electrode grids, meaning that all available electrodes were used, was 75±17% across participants using HFB, 22±5% using only the beta band activity, and 54±17% for the beta band and HFB combined. Although these results were significantly above the practical chance level of 16.7%, there was considerable variability between participants. Interestingly, this variability was not significantly correlated to the number of electrodes per participant (p = 0.07, 0.17 and 0.10 for HFB, beta and HFB/beta). Since we sought to maximize classification performance, further analyses were performed using only HFB features.

### 3.3 Subgrid analysis

We aimed to determine whether small electrode grids could achieve a comparable level of classification accuracy to that of the full 64-128 electrode grids. As detailed in Section 2.6, we performed an exhaustive grid search of all possible rectangular subgrids for each electrode grid, comparing their performance in word classification accuracy. Across participants, 70% of the top ten best-performing subgrids contained 32 electrodes or less. On the graph of cumulative classification accuracy, 32 electrodes corresponded to an elbow, with sharp increases in accuracy for subgrids of 1 to 32 electrodes and gradual trail-off in accuracy for subgrids larger than 32 electrodes (Figure 5).

**Figure 5.**
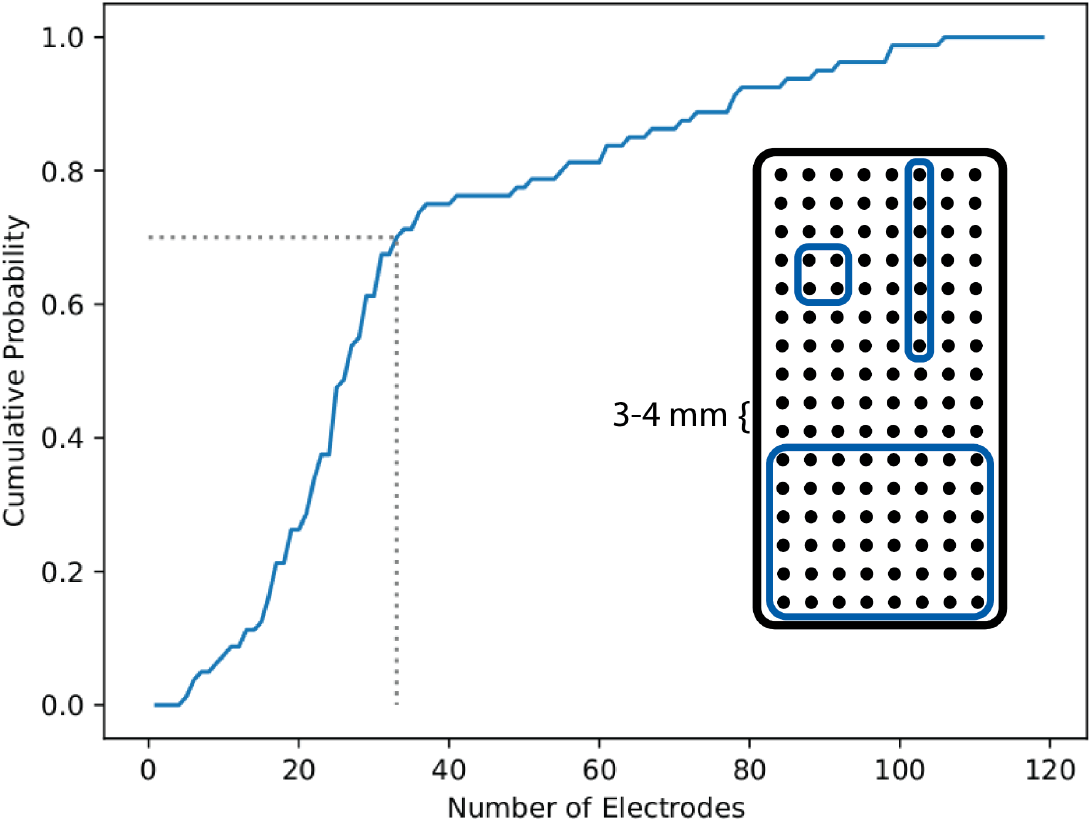
Cumulative distribution of best-performing subgrid sizes by electrode count. All possible rectangular subgrids were evaluated, with illustrated examples including a 2×2, a 1×7, and an 8×6-electrode grid. For this figure, the 10 subgrids with the highest classification accuracy for each participant were selected, and their size distribution is shown.

Based on this result, we further investigated results for 32-electrode subgrids. Compared to the original grids, this represented a 50 to 80% reduction in grid sizes, with the full grids covering areas of 1,225 mm^2^ to 2,806 mm^2^, and the subgrids covering areas from 325 mm^2^ to 561 mm^2^.

For each participant, classification was performed on every possible 4×8 electrode subgrid, oriented both vertically and horizontally. The median classification accuracy of the best-performing subgrids across cross-validation test sets was 76±16%, while the median accuracy using full grids was 75±17%. These accuracies did not differ significantly (Z_full-subgrids_= -1.2, p = 0.22, Wilcoxon signed-rank test), suggesting that a smaller, 32-electrode grid, if placed optimally, could reach the same level of performance as a larger electrode grid for the classification of twelve different words. Adjusting the common average reference to only include electrodes in smaller subgrids did not lead to significantly different results (77±20%, Z_full-subgrid_= 5.0, p = 0.49, Wilcoxon signed-rank test).

Across participants, the optimal location of the 32-electrode subgrids stayed consistent across cross-validation test folds (Figure 6). Notably, the best-performing subgrids were consistently arranged vertically on the SMC, and were frequently positioned to cover the bulk of the electrodes identified as significant in the t-test comparing speech vs silence (Section 3.1). Participants with the least consistent subgrid placements over test folds either had a grid with a larger electrode diameter and inter-electrode distance compared to others (P1) or had a horizontally positioned grid (P8).

**Figure 6.**
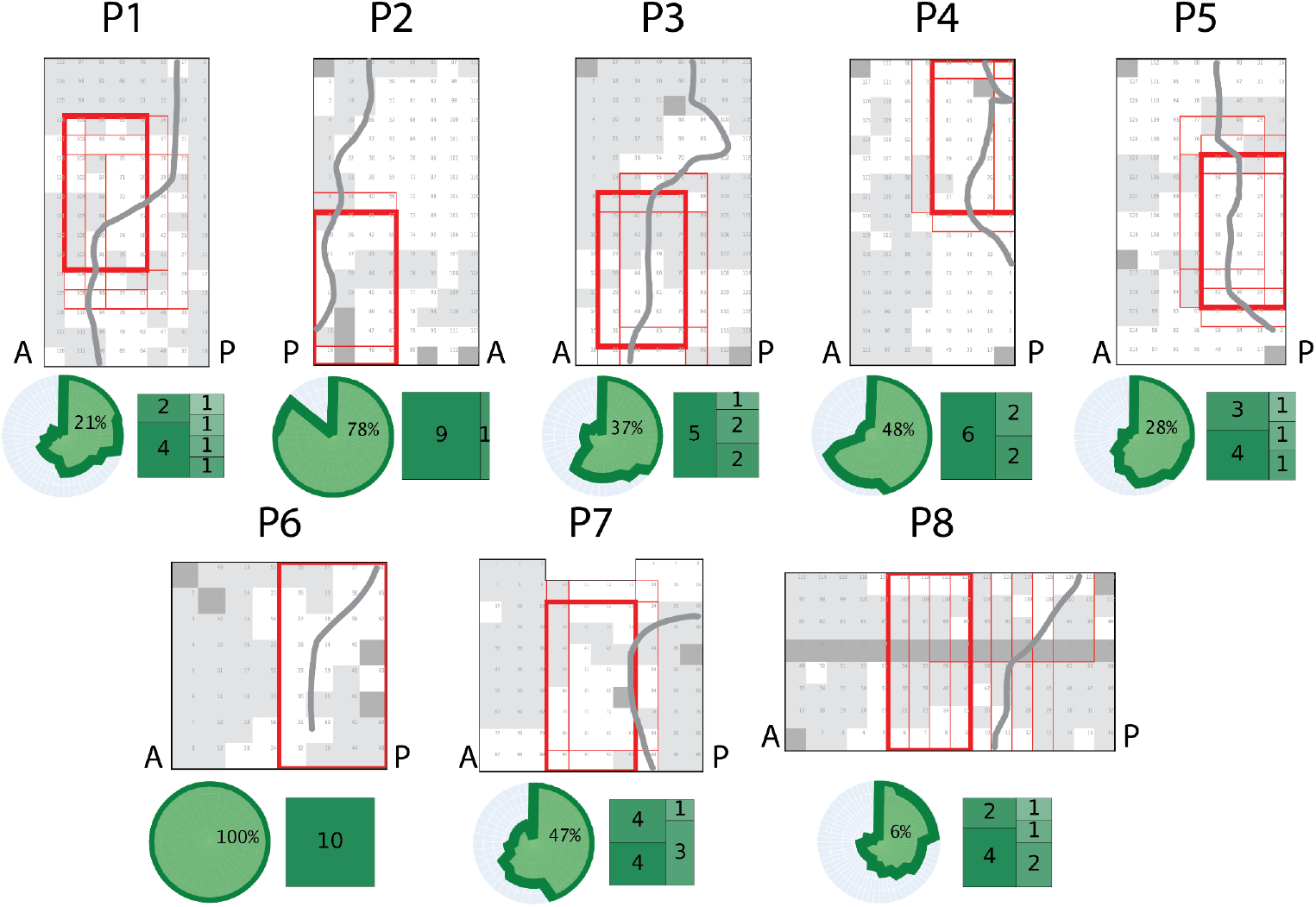
Best 32-electrode subgrids per participant. For each participant, the optimal subgrids selected across cross-validation folds are shown in red. White squares denote electrodes with significant speech vs. silence differences in HFB power based on the t-test, while light gray squares denote electrodes with insignificant differences. Dark gray squares denote electrodes that were excluded from analyses (see Methods). The most frequently selected subgrid with the best performance across folds is highlighted in bold red, and central sulci are drawn in dark gray. In participant P7, two subgrids were chosen in four folds each; however, to maintain consistency across participants, only one was highlighted and used in subsequent analyses, as both grids had nearly identical performance. Below each electrode grid, a radial plot illustrates the overlap of the top 32-electrode subgrids across folds. Electrodes are plotted on the angular axis, with the radial axis indicating how many folds they were chosen in. The percentage reflects the proportion of electrodes selected in every fold; for example, in participant P2, 78% of electrodes were consistently chosen in all folds, with the remaining electrodes rarely selected, as represented by the deep insertion in the circle. In contrast, in participant P1, only 21% of electrodes were chosen in all folds, with the remaining electrodes being chosen in a varying number of folds. The tree plots display the distribution of best-performing subgrids across folds, where each cell represents a unique subgrid and its frequency across folds. As an example, in P1, the most common subgrid was chosen 4 times, the second most common subgrid was chosen twice, and all other subgrids performed best only in one cross-validation fold. This analysis used only HFB features.

Mapping the best-performing 32-electrode subgrid for each participant onto the common MNI space revealed a considerable positional overlap across participants (Figure 7). It is also notable that the optimal subgrids are clustered around the central sulcus, covering both pre- and postcentral areas.

**Figure 7.**
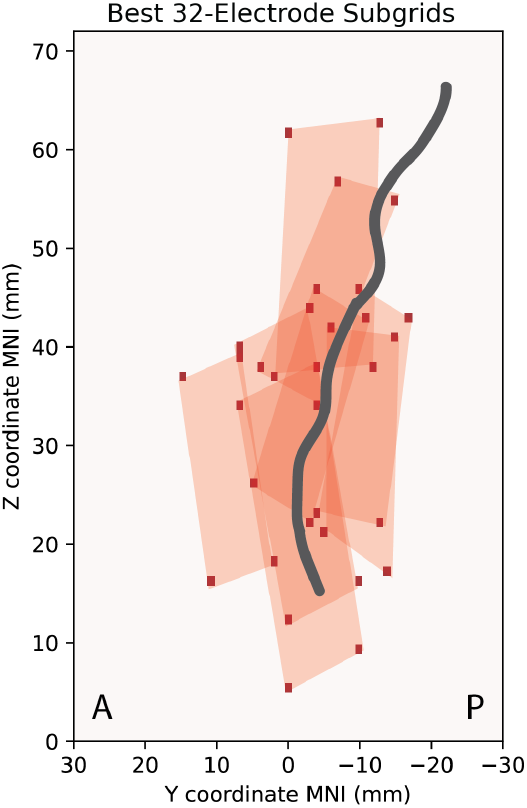
The best 32-electrode subgrid for each participant mapped onto the common MNI space. The best-performing subgrid for each participant is the one that was found most frequently across cross-validation folds. Points in dark red outline corner electrodes of each participant’s subgrid. The central sulcus is drawn in dark gray. Only HFB features were used in this analysis. For visualization purposes, the subgrid of P2 is mapped onto the left hemisphere.

### 3.4 Recursive feature elimination

To identify the most informative electrode locations for classification without the 4×8 configuration constraint, we applied recursive feature elimination (RFE) to the full set of electrodes, focusing exclusively on HFB features. To enable a fair comparison with the previous analysis, the optimal result was defined as the RFE step that achieved the highest accuracy using no more than 32 electrodes.

Across participants, RFE identified that a median of 23±8 electrodes was needed to achieve the best classification accuracy (Figure S1). The median accuracy reached 80±20%, and did not differ significantly from the accuracy reached with 32-electrode subgrids (76±16%, Z_RFE-grids_= -1.0, p = 0.33 Wilcoxon signed-rank test). The difference between RFE and full grids did not reach significance when Bonferroni corrected for multiple comparisons (75±17%, Z_RFE-full_= -2.1, p = 0.04 Wilcoxon signed-rank test).

Interestingly, electrodes selected by the RFE procedure did not entirely overlap with the previously identified foci of speech-related activity (Figure 8, S3). Across participants, an average of about 25% of the RFE-selected electrodes were not significant in the t-test. In addition, there was no complete overlap with the best-performing subgrids of 32 electrodes (Figure 9, S4).

**Figure 8.**
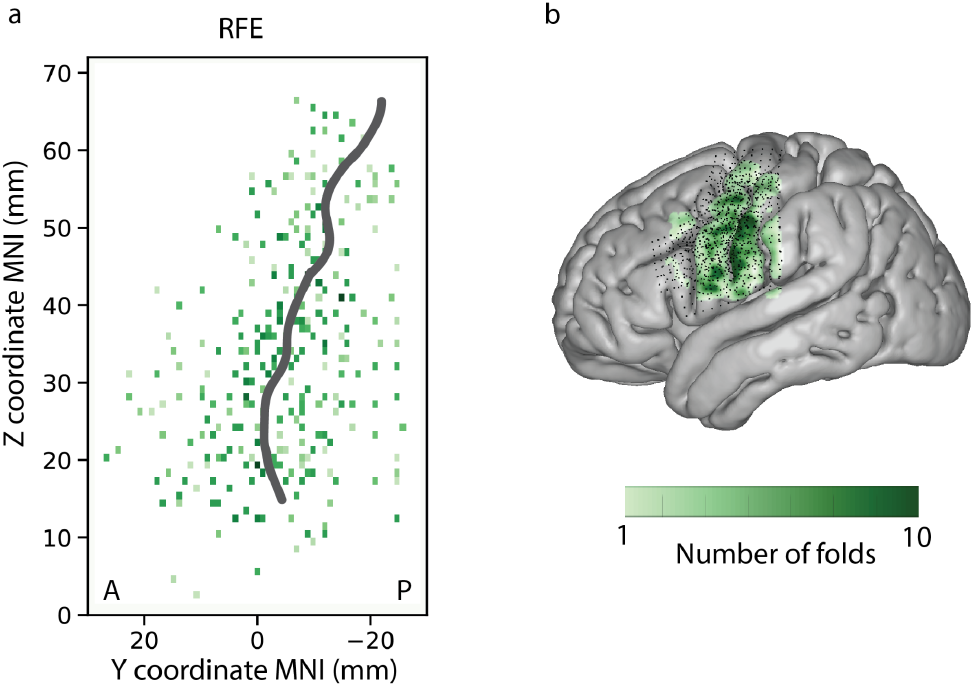
Number of folds in which each electrode was selected in the RFE algorithm, mapped onto the common MNI space. a) MNI coordinates in a 2D space (x-coordinate removed), with anterior and posterior directions labeled as “A” and “P”. b) Results plotted on the MNI brain, with the black dots corresponding to all individual electrodes. In both subfigures, data from P2 is mapped onto the left hemisphere (LH).

**Figure 9.**
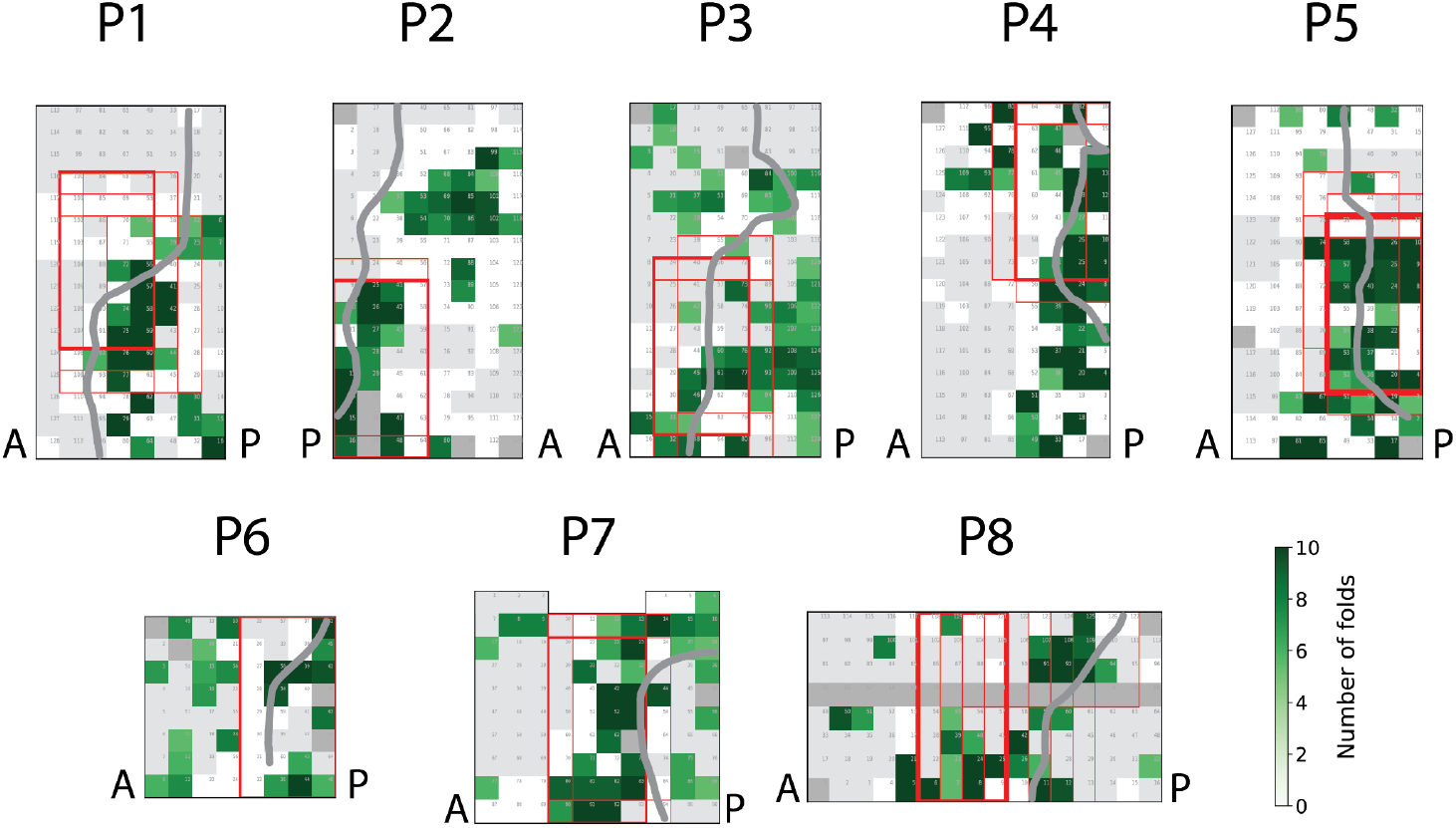
Best electrodes as determined by subgrid analysis and RFE. Indicated in green are electrodes determined by the RFE algorithm to be most informative – darker green means they were chosen in more cross-validation folds. Electrodes in light gray showed no significant difference between speech and silence, as revealed by the t-test in 3.1, and were not identified in the RFE analysis. Electrodes in dark gray were removed from all analyses due to having noisy signals or being reference electrodes. Individual 32-electrode subgrids across 10 cross-validation test folds are shown in red. The most frequently found 32-electrode subgrid with best performance across folds is in bold red. The central sulci are drawn in dark gray. “A” and “P” denote anterior and posterior directions on the cortex of participants.

### 3.5 Recursive feature elimination on preselected electrodes

As a final analysis, we repeated the RFE procedure while restricting the RFE’s initial feature space only to those electrodes that were significant in the t-test comparing speech and silence fragments (t-test preselection). We then compared the classification accuracy obtained as such with the classification accuracy of the original RFE (no preselection) (Figure 10). We found no significant difference in classification accuracy between the RFE procedures with (77±21%) and without electrode preselection (80±20%, Z_preselected-all_= -0.1, p = 0.92 Wilcoxon signed-rank test), implying that the information necessary to distinguish words can be extracted from electrodes that showed a significant increase in activity during speech compared to silence.

**Figure 10.**
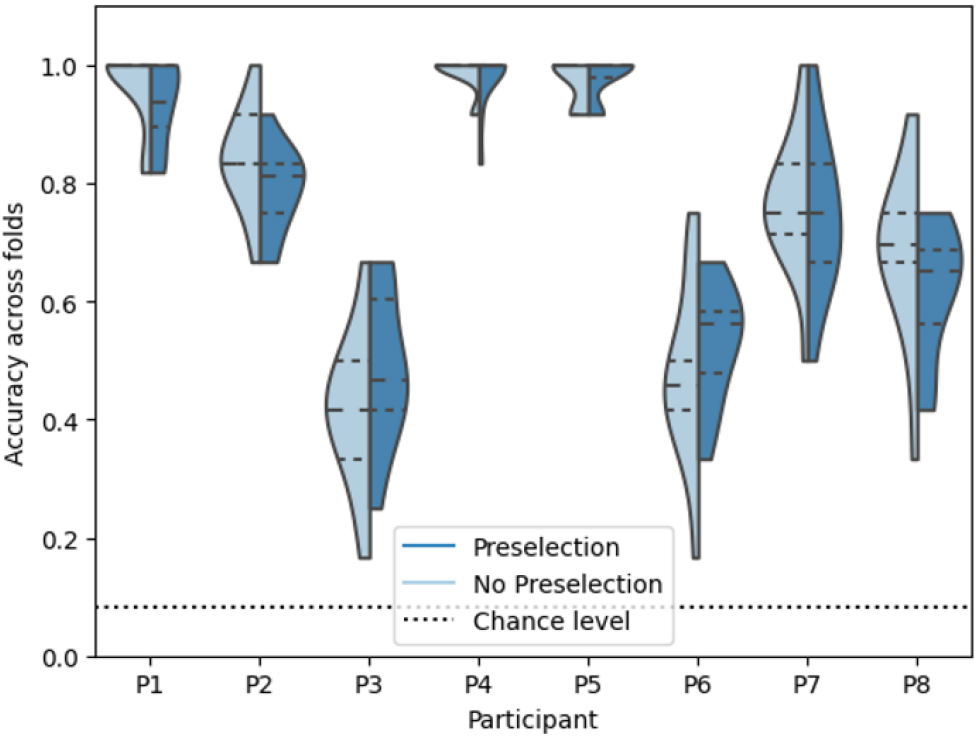
Classification accuracy across cross-validation test folds for each participant with RFE on HFB features, with no electrode preselection (light blue) and with electrode preselection based on the t-test comparing speech with silent fragments (dark blue). The dashed lines show the 25th, 50th and 75th quartiles of the data, and the density shows the full data range.

## 4. Discussion

The present study investigated the feasibility of using optimally positioned HD-ECoG grids of limited size for word classification in BCIs. Our findings show that configurations of only 32 electrodes with surface areas of 325 mm^2^ to 561 mm^2^ can achieve classification accuracy comparable to larger grids of 64 to 128 electrodes spanning large parts of the SMC (1,225 mm^2^ to 2,806 mm^2^ surface area). These smaller grids performed best when oriented vertically and centered on the central sulcus. Additionally, we identified two distinct SMC foci (one in ventral and one in dorsal precentral SMC) of HFB activity during speech. These regions exhibited a significant response to the word repetition task, and also encoded word-specific information as shown by a recursive feature elimination analysis confined to the speech-significant areas.

Smaller ECoG grids may be more desirable for BCI since they can be implanted with a lower surgical burden as well as smaller risks of complications and infections (Hamer et al., 2002). One study on grids used for epilepsy monitoring (interelectrode distance of 1 cm, median implantation duration of 11 days) showed that the size of implanted grids was the sole independent predictor of increased complications following implantation, with grids of size 4×8 or smaller significantly reducing complications compared to larger grids (Rahman et al., 2016).

Additionally, smaller grids may be implanted with even lower surgical burden in the near future. Recently, a technique was developed for implanting ECoG grids with an area of 156 mm^2^ through a cranial micro-slit (Hettick et al., 2022), and another method using expandable grids allowed for implantation of a 32-electrode ECoG grid with a coverage of 600 mm^2^ through a 12mm burr-hole (Coles et al., 2024). Both techniques eliminate the need for a craniotomy, significantly reducing the complexity and potential complications during grid implantation and explantation procedures.

Limited previous work has attempted to determine the minimal size of ECoG grids needed for high classification accuracy. For example, a simulation study using high resolution functional magnetic resonance imaging (fMRI) found that accurate classification of three hand gestures from SMC can be achieved with 3×3 electrodes and an inter-electrode distance of 8 mm (Van Den Boom et al., 2021), having a total area of 361 mm^2^. In another study, areas of less than 100 mm^2^ were sufficient for classifying four articulator movements with HD-ECoG (Salari et al., 2019). The present findings extend this previous work to a larger number of classes (twelve words) and apply this work to classification of speech from SMC.

We propose that smaller ECoG grid configurations may be used for word classification BCIs without compromising performance, but this requires specific grid positioning. Based on our results, we conclude that such smaller grids should be positioned over the central sulcus, covering both precentral and postcentral areas, as top-performing configurations in our study were oriented vertically and clustered around the central sulcus in all participants. We speculate that the vertical positioning of the optimal configurations might enable better integration of information from the two foci of HFB activity we observed during speech, one in the dorsal and one in the ventral precentral SMC. Anatomically, the dorsal area of high HFB activity was located near the midpoint of the precentral gyrus. This area, called the middle precentral gyrus or area 55b, has been associated with important functions relevant for speech processing, and is thought to be an important integration cortical hub for both dorsal and ventral streams of language (Glasser et al., 2016, Hazem et al., 2021, Silva et al., 2022). Previous work has demonstrated successful speech decoding in an individual with ALS based on brain activity in area 55b (Card et al., 2024). The second area showing strong HFB activity during speech overlapped with an area of interest in the dorsal part of the ventral premotor cortex (6v in the Human Connectome Project). This area is implicated in both language production and processing, and has yielded good results for speech BCIs (Glasser et al., 2016, Willett et al., 2023). Both areas are thus known to be of value for speech decoding and word classification. As shown in Figure 4, the foci of HFB activity found during speech partially overlapped with areas 55b and 6v. However, the regions of peak activation appeared slightly more dorsal than the previously found areas, likely reflecting inter-individual variability across the eight participants. Additionally, when mapping areas 55b and 6v back to individual space, we found that optimal subgrids overlapped with both areas whenever possible (see Figure S5). However, in all participants, these optimal subgrids were located more posteriorly than areas 55b and 6v and were generally centered around the central sulcus, therefore also covering parts of the postcentral gyrus.

RFE does not yield a contiguous subgrid, making it less useful for guiding minimally invasive implantation; however, it can still highlight the most informative electrodes for word classification. The electrodes selected by the unconstrained RFE procedure were not located exclusively in the two foci of activity discussed in the previous paragraph. This spatial variability, paired with similar classification accuracy using full grids, subgrids and RFE suggests redundancy in the cortical patterns evoked by speech — the information necessary for classifying words is available across multiple, or relatively extensive, SMC areas. Interestingly, when constrained to only the significantly activated electrodes (Section 3.5), RFE still reached the same level of accuracy as when applied to the full grids. This suggests that electrodes sensitive to speech as opposed to silence also contain word-discriminative information. Supporting this, only a small number of electrodes selected in more than half of the folds were not significantly activated by speech (Figure S4). Thus, while redundant information exists, the consistently chosen electrodes in the RFE analysis tended to fall within the regions with significant HFB activity, in line with the optimal positioning of subgrids over these areas. Information that is critical for implantation can thus be gained by comparing HFB between speech and silence. To conduct such a test, non-invasive methods such as fMRI could be used, since the blood-oxygen dependent level measured with fMRI is highly correlated with HFB (Lachaux et al., 2007, Hermes et al., 2012, Siero et al., 2014).

Additionally, RFE indicated that electrodes providing most information for word classification were located not only in precentral gyrus, but also the postcentral gyrus. This finding is in line with the positioning of the most optimal subgrids over the central sulcus, and underscores the contribution of the somatosensory cortex to decoding. Further work is needed to explore the role of both the precentral areas like 55b and 6v, as well as the postcentral gyrus and somatosensory processing in speech production, not only in able-bodied individuals but also in people who cannot move their facial muscles, the target users of speech BCIs. Since people with paralysis extending to the face cannot physically perform the attempted movements, activity in the somatosensory cortex will likely be less prominent. Recent findings, however, indicate that the primary somatosensory cortex receives motor output information before the arrival of sensory feedback signals (Sun et al., 2015, Umeda et al., 2019), contributing to theories of motor control that involve feedforward and efference copy concepts (Wolpert & Kawato, 1998).

Successful motor decoding from the somatosensory cortex has also been achieved both in amputees (Bruurmijn et al., 2017) and in people with locked-in syndrome due to ALS (Leinders et al., 2023), encouraging generalizability of the results to people with severe paralysis.

There are several limitations to the present study. First, the number of participants was limited, and the classification performance between participants varied considerably, between 40 and 100% accuracy. This could be due to differences in signal quality, participant alertness, task performance, medication and epileptic activity, despite the grids being placed on presumed healthy tissue. Importantly, full grid classification accuracy was not correlated with original grid size. Additionally, two participants (P4 and P5) showed increases in the beta and alpha band power during speech, contrary to the expected decrease, and several participants showed decreases in the HFB power in a small number of electrodes (Figure 3). These activity changes are unlikely to be associated with noisy electrodes or the common average reference, leaving the origin of these deviations unclear. Despite these unexplained differences, however, our main findings remained consistent across all participants.

Second, the current study involved a closed set of twelve words. Expanding the vocabulary and improving continuous speech decoding, in particular with the aid of language models, has become the focus of the latest speech BCI research. It is unclear how using smaller, optimally positioned grid configurations would affect continuous speech decoding. However, classifying a limited set of words—such as for controlling a computer menu or a smart home with a few key commands—likely serves as a valuable standalone BCI-based communication method. This approach may be especially useful depending on user preferences and situational needs, given uncertainties about speech decodability in anarthria and the challenges of developing high-electrode-count implants.

Future work is needed to reproduce these findings in people with severe paralysis – potential users that stand to benefit from speech BCI technology the most. It would also be of interest to see how these results would scale with the level of task complexity, and whether larger grid coverage may be necessary to reach high classification accuracy for larger sets of words. Finally, more work is needed to explore the role of electrode density in the context of speech classification and determine whether higher-density grids could enable even smaller coverage areas.

## 5. Conclusion

Our study demonstrates that a small ECoG grid of 32 electrodes with a cortical area of 325 mm^2^ to 561 mm^2^, if positioned optimally, can match the word classification accuracy of larger grids, thus paving the way for less invasive and more efficient speech BCIs. By identifying critical regions within the SMC and showing the predictive value of speech to silence contrasts, we offer a practical approach to optimizing small grid placement and feature selection. These findings can further the development of speech BCIs for individuals with severe paralysis offering them a minimally invasive yet effective means of communication.

## Acknowledgements

This publication is part of the project Dutch Brain Interface Initiative (DBI2) with project number 024.005.022 of the research programme Gravitation, which is financed by the Dutch Ministry of Education, Culture, and Science (OCW) via the Dutch Research Council (NWO). In addition, this project is funded by the European Union’s HORIZON-EIC-2021-PATHFINDER CHALLENGES program under grant agreement No 101070939 and by the Swiss State Secretariat for Education, Research and Innovation (SERI) under contract number 22.00198.

