## Supplementary Figures for "Optimal positioning and size of high-density electrocorticography grids for speech brain-computer interfaces"

**Supplementary Material**

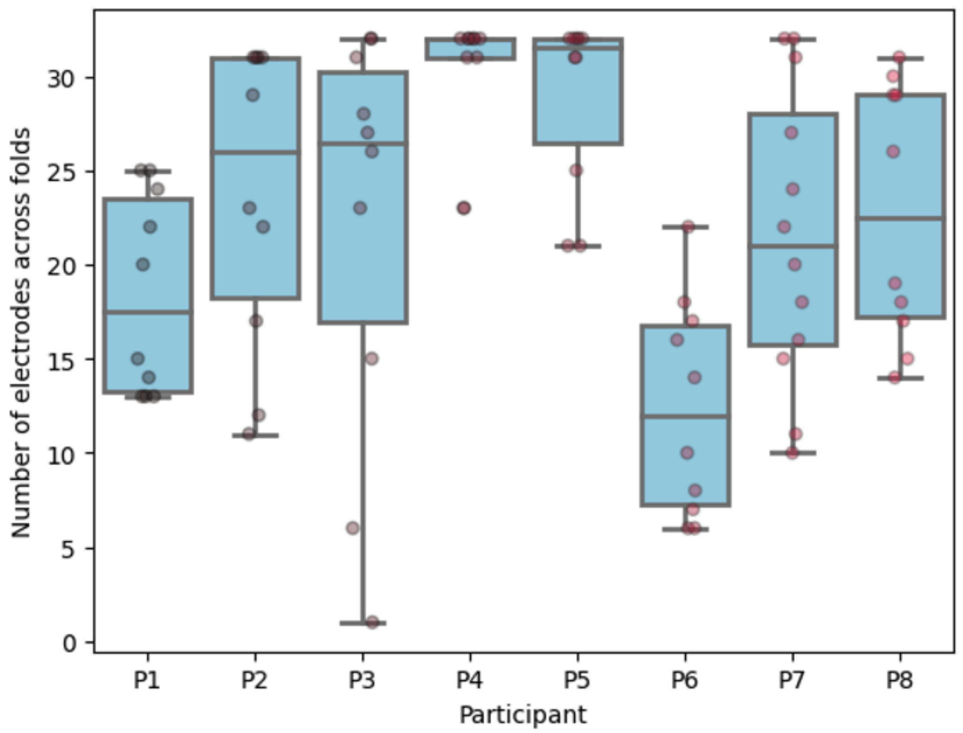


Figure S1: Number of electrodes chosen for each participant across the 10 cross-validation folds by the RFE algorithm, which was capped at 32 electrodes for better comparison to the subgrid analysis as described in Section 3.3.

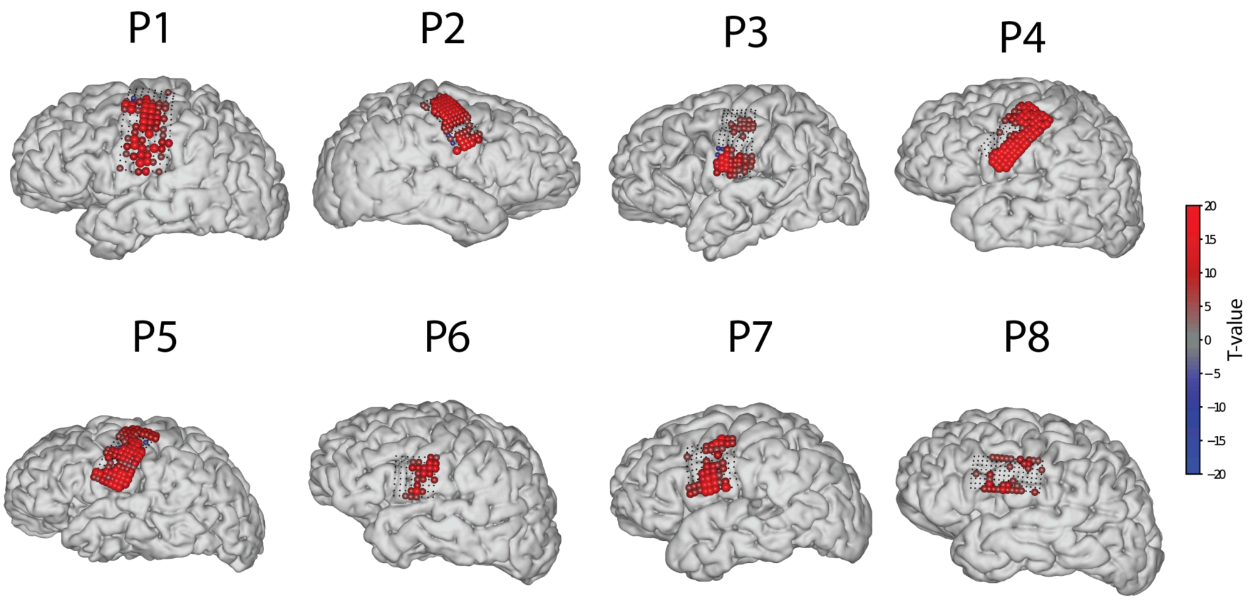


Figure S2: The electrodes with a significant difference in mean HFB activity during silence and speech fragments, displayed on the individual brains of the participants.

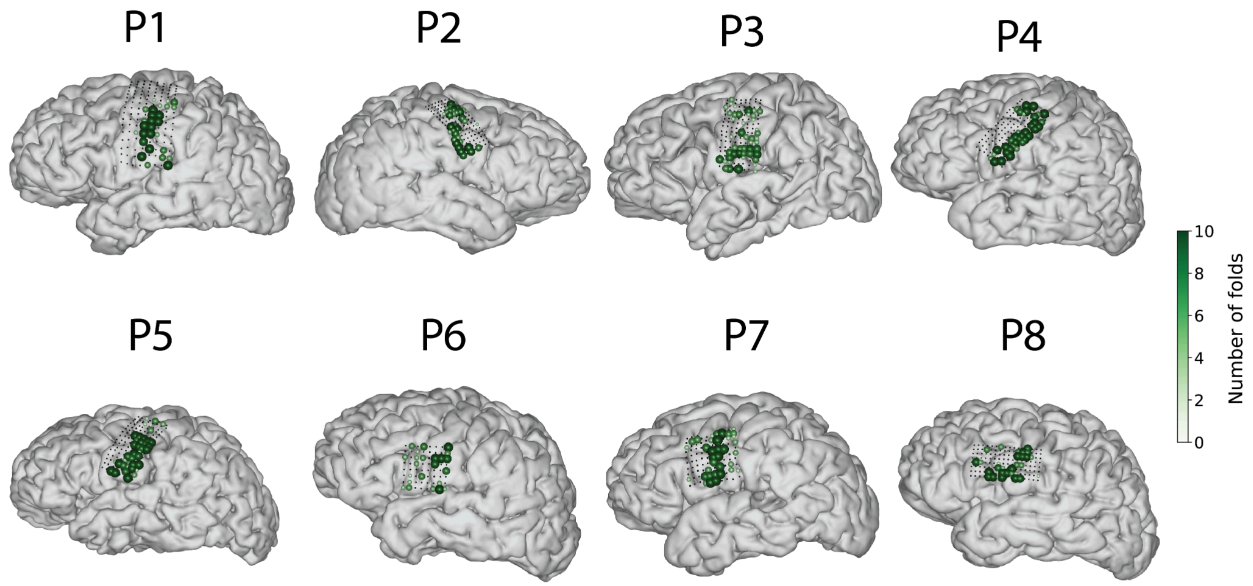


Figure S3: The electrodes chosen by the RFE-algorithm as described in Section 2.7, with word classification using HFB activity, displayed on the individual brains of the participants. Darker electrodes were chosen more frequently across cross-validation folds.

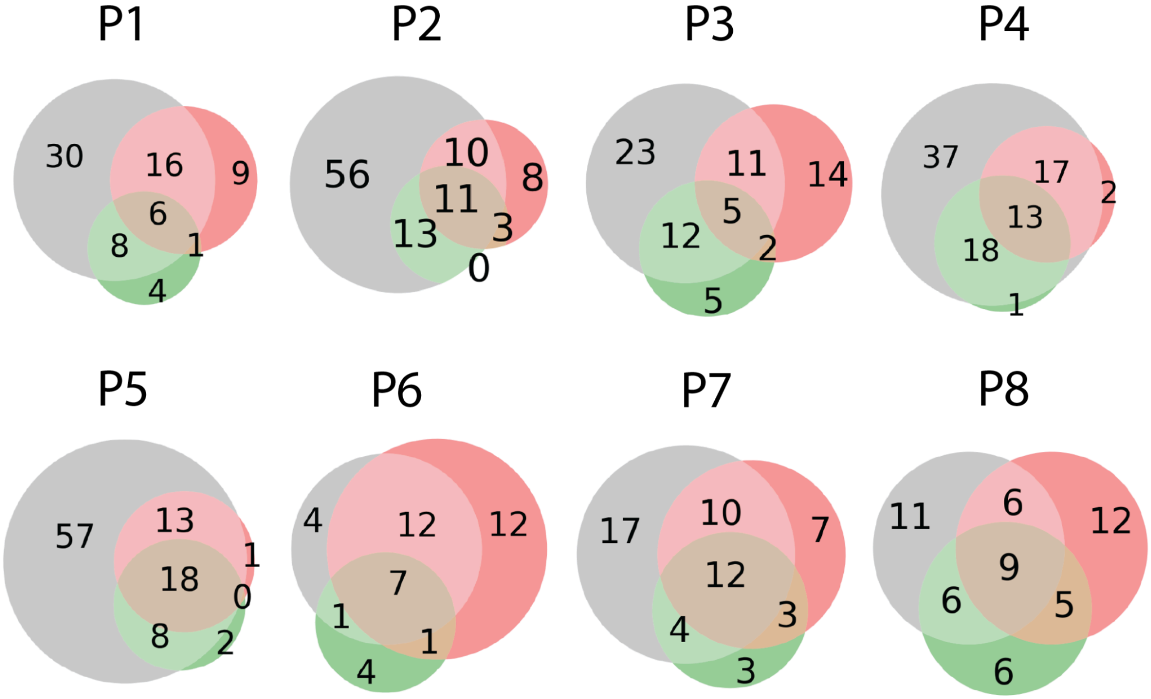


Figure S4: Overlap between electrodes with a significant difference in speech and silence (gray), electrodes in the best-performing subgrid across cross-validation folds (red), and chosen by the RFE-algorithm in 50% or more of all folds (green) for each participant.

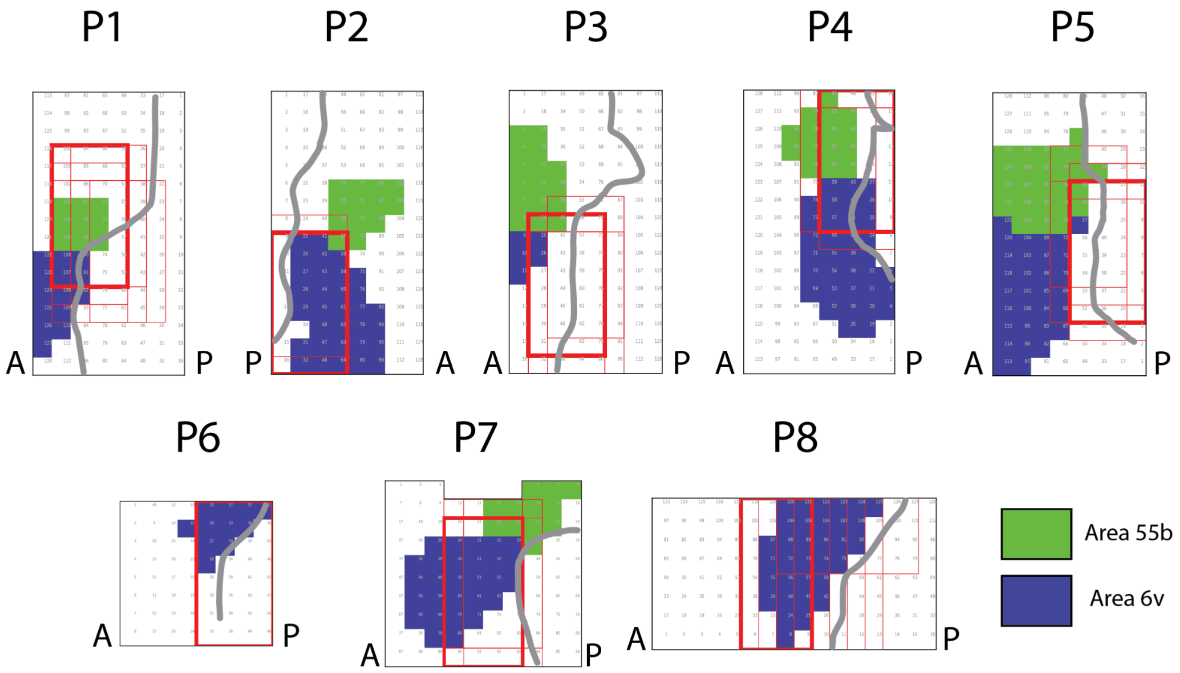


Figure S5: Areas 55b and 6v with best-performing subgrid per cross-validation fold. Electrodes are marked in green if their MNI-space distance to any point in area 55b was less than 5 mm, and in blue if they met the same criterion for area 6v.
